# TMS–EEG Indices to Define Local Cortical Excitability Thresholds

**DOI:** 10.64898/2026.01.12.699003

**Authors:** Ilkka Rissanen, Ida Granö, Victor Souza, Sabin Sathyan, Nivethida Thirugnanasambandam, Risto J. Ilmoniemi, Pantelis Lioumis

## Abstract

**Introduction:** Transcranial magnetic stimulation (TMS) is widely employed to treat various psychiatric and neurological disorders. However, TMS protocols typically rely on generalizations, particularly in selecting stimulation intensities, leading to suboptimal and variable outcomes. Combining TMS with electroencephalography (EEG) offers a potential solution by allowing direct monitoring of stimulation effects. In this study, we investigate how features of the TMS–EEG signal change with intensity to identify thresholds implying qualitative shifts in the brain response.

**Methods:** We stimulated eight subjects at both the primary motor cortex (M1) and the pre-supplementary motor area (pre-SMA) with navigated TMS at 15 closely spaced intensities and measured TMS-evoked EEG responses (TMS-evoked potential, TEP) with 60 trials per intensity. TEP thresholds were identified with three methods: by selecting the intensity where the TEP peak-to-peak exceeds 6 μV, or by fitting either piecewise-linear or sigmoid curves into the power spectral density (PSD) frequency components to identify nonlinear intensity behavior.

**Results:** The identified TEP thresholds varied depending on subject, target, and identification method. In M1, the thresholds attained with the fixed-amplitude and piecewise PSD fit methods averaged around the motor threshold, and in pre-SMA around 120% of the motor threshold. The TEP thresholds yielded by the sigmoid fit method were higher in intensity, and least consistent between subjects.

**Conclusions:** Our findings support the hypothesis that detectable changes in EEG patterns occur at specific TMS intensities. These results provide a basis for individualized stimulation dosing, potentially enhancing therapeutic efficacy and reliability. Future research should focus on refining these methods and validating their clinical applicability across diverse conditions and patient populations.

## Introduction

Transcranial magnetic stimulation (TMS) is a noninvasive brain stimulation method that activates cortical neurons with magnetic pulses (Barker et al., 1985). It is widely used for treating depression and neurological conditions such as chronic pain and movement disorders, alongside functional cortical motor and speech mapping for presurgical evaluation (Hui et al., 2020; Lefaucheur et al., 2020; Rossi et al., 2021; Trapp et al., 2025). However, contemporary clinical TMS protocols rely on broad generalizations when selecting stimulation intensities (Lefaucheur et al., 2020; Platz, 2016). Such lack of individualization is likely to result in variable and suboptimal treatment outcomes, a common phenomenon reported in clinical trials (Berlim et al., 2014; Chang et al., 2024). To enable individualized selection of stimulation intensity based on subject and stimulation target for optimal therapeutic effect, we need a better understanding of the relation between stimulation intensity and brain response. The combination of TMS with electroencephalography (EEG) enables the recording of cortical reactivity to the stimulation and its propagation to other brain regions (Hernandez-Pavon et al., 2023; Ilmoniemi et al., 1997; Ilmoniemi & Kičić, 2010; Massimini et al., 2005; Parmigiani et al., 2023; Wischnewski et al., 2024), thereby paving the way for personalized dosing tailored to individual subjects (Casarotto et al., 2022; Lioumis & Rosanova, 2022; Parmigiani et al., 2025; Ukharova et al., 2025).

The most common approach to determine stimulation intensity for non-motor brain regions is based on the motor threshold (MT), defined as the minimum intensity required to consistently elicit a specific muscle response when stimulating the primary motor cortex (M1) (Rossi et al., 2021). For example, prominent TMS treatment protocols for depression stimulate at 120% of MT at the left dorsolateral prefrontal cortex (Lefaucheur et al., 2020; Trapp et al., 2025). However, as responsiveness to stimulation has subject-dependent differences between brain regions (Ozdemir et al., 2021; Rosanova et al., 2009), for some patients, sufficient stimulation intensity outside the motor cortex can differ significantly from the MT (Harquel et al., 2016; Trapp et al., 2025), making it an unreliable reference value for most therapeutic TMS protocols.

The combination of EEG with TMS enables empirical determination of the appropriate stimulation intensity and orientation directly from the induced, subject-specific brain activity (Casarotto et al., 2022; Tervo et al., 2022). However, merely determining the minimum intensity required to excite cortical neurons is likely not sufficient. In the motor cortex, pulses can induce neuronal perturbation visible in EEG even if they are not strong enough to elicit a muscle response (Kähkönen et al., 2005; Komssi et al., 2004). Likewise, in other brain regions, TMS pulses that induce only minor cortical activation may not engage the networks of the brain to achieve desired therapeutic outcomes. It is essential to determine the threshold of stimulation intensity that yields a functionally relevant causal response, rather than only a transient, local disturbance. Such thresholds could be identified by observing qualitative changes with respect to stimulation intensity in EEG responses at the stimulation target. These differences in TEP shape could be extracted from frequency-domain analysis, as frequency spectra characterize the overall shape of the response. Distinct oscillatory patterns have been found to be related to function-specific brain circuits (Lioumis et al., 2025; Rosanova et al., 2009), and previous works have investigated characterizing TMS–EEG responses in the frequency domain (Raffin et al., 2020; Saari et al., 2018).

Previous research on the effect of stimulation intensity on EEG response has generally used broad and sparse ranges of intensities. For example, Kähkönen et al. (2005) and Komssi et al. (2004) investigated the effect of TMS intensity in the M1 and left middle frontal gyrus with four intensities: 60, 80, 100, and 120% of resting MT (rMT). They reported linear or early-sigmoid behavior for examined peaks in the TMS-evoked potential (TEP). Casali and colleagues (Casali et al., 2010) demonstrated that increasing intensity made TMS-evoked activation last longer with a wider spread of activity and identified an intensity threshold of 40–50 V/m in the cortex to elicit brain activity visible in EEG. On the other hand, Komssi et al. (2007) identified 33–44 V/m as the threshold to elicit minimal cortical activation, while demonstrating that the presence of various peaks in the TEP depends on stimulation intensity. However, a high-resolution empirical model for describing how TEP parameters change at varying stimulus intensities is still missing. In particular, we lack the means to identify TEP-based intensity thresholds that indicate a qualitative shift in brain response (Casarotto et al., 2022; Lioumis & Rosanova, 2022).

The aim of this research was to identify a distinct alteration in cortical reactivity corresponding to a threshold-level neural response. As evidenced by overt motor responses only occurring with sufficient stimulation intensity, we hypothesized that there would be patterns of TMS-induced brain activation that only occur above a certain intensity, and that crossing such thresholds would produce a detectable change in EEG behavior. We developed a frequency-domain-based method for determining TEP intensity thresholds and compared the results to those of a method based on deflection amplitudes.

## Methods

### Participants

Eight healthy volunteers free from neurological and motor deficits participated in this study (ages 25–51 years, right-handed, 5 male). Each participant underwent two experimental sessions on different days: one stimulating M1, the other stimulating pre-SMA. The experimental protocols were approved by the Coordinating Ethics Committee of Helsinki University Hospital. All subjects signed a written informed consent.

### Experimental procedure

Stimulation was performed using a navigated TMS system (NBT 5.0, Nexstim Plc., Finland) with a figure-of-eight coil (70-mm radius; Cooled Coil, Nexstim) and individual T1-weighted magnetic resonance images (MRIs). The MRIs were acquired with a volumetric gradient echo sequence (1 x 1 x 1 mm^3^ voxel size, 240 x 240 x 240 acquisition matrix) with repetition and echo times of 2.53 and 3.3 ms, respectively, in a Skyra 3T scanner (Siemens Healthcare, Germany). EEG signals were recorded with a BrainAMP EEG system (Brain Products, GmbH, Germany) and a 62-channel EEG cap (Easycap) with passive Ag/AgCl-sintered multitrode electrodes. The EEG signals were low-pass filtered at 1000 Hz and sampled at 5000 Hz. When targeting M1, electromyography (EMG) was measured with a Nexstim EMG system with electrodes placed in a belly–tendon montage on the right *abductor pollicis brevis* (APB) muscle. During measurements, active noise masking consisting of white noise mixed with recorded coil clicks was played to the subject through inserted earphones (ER3C Insert Earphones, Etymotic Research Inc., United States) to minimize auditory-evoked responses (Russo et al., 2022; https://github.com/iTCf/TAAC).

Each measurement session consisted of four steps: *i*) EEG and noise masking preparations, *ii*) determining the stimulation target, *iii*) determining stimulation reference intensity, and *iv*) recording TEPs at the selected target with varying intensities.

EEG electrode contacts were prepared by scraping the scalp under the electrodes with conductive abrasive paste (OneStep AbrasivPlus, H + H Medical Devices, Germany), after which the electrodes were filled with conductive gel (Electro-Gel, ECI, Netherlands). Electrode impedances were kept below 5 kΩ; they were periodically checked during the experiment and adjusted if their impedance had increased. The reference electrode was placed on the right mastoid and the ground electrode on the right zygomatic bone. The noise masking volume was adjusted until the subject reported being unable to perceive the coil clicks produced by the TMS coil held 5–10 cm above the vertex, stimulating at 80% maximum stimulator output (MSO). The absence of auditory responses was confirmed by visual inspection of 30 trial-averaged EEG responses from a real-time EEG visualization tool (Casarotto et al., 2022).

After preparations, the subject was seated in a comfortable chair and instructed to stay relaxed and look at a black cross 2–3 meters away. When targeting M1, the APB hotspot was used as the target, determined by finding the coil location and orientation that produced consistent and maximal motor-evoked potentials (MEPs) at a suprathreshold intensity. The rMT of APB was estimated by the Nexstim system’s built-in maximum-likelihood algorithm (Awiszus, 2003). If APB hotspot stimulation caused significant muscle artifact on EEG, the optimal stimulation target was determined by moving and rotating the coil within M1 to find the coil position that produced artifact-free TEPs with early (<50 ms) peak-to-peak amplitudes of 6–10 µV in electrodes under the stimulation coil (Casarotto et al., 2022) at minimal stimulation intensity. If no site fulfilled the above conditions, the target with the smallest artifacts that yielded a visible TEP was chosen. When targeting pre-SMA, we performed the search for artifact-free TEPs in the area, as identified by individual anatomical landmarks (Ferrarelli et al., 2008; Rosanova et al., 2009). The coil orientations were generally perpendicular to the targeted sulci, and thus approximately perpendicular to the central sulcus for M1 stimulation, and latero–medial for pre-SMA stimulation.

Having defined the target, we determined the reference intensity by finding the highest stimulator output with no visible TEPs in the averaged 20 TMS–EEG responses (Casali et al., 2010). Starting from the reference level, we used 15 intensities at increments of 2% MSO for the measurements. Then, 20 TMS–EEG responses were measured at the highest stimulation intensity to confirm the lack of scalp-muscle activations. If significant activation occurred, the stimulation target and starting intensity were readjusted. If the highest intensity exceeded 80% MSO, the noise masking volume was increased to ensure the absence of auditory-evoked responses.

As the last step, we recorded TEPs at the 15 chosen intensities. A block of stimuli consisting of 60 pulses was delivered at each intensity at an interstimulus interval jittered between 2.0 and 2.3 s. The sequence of blocks was pseudo-randomized. The coil position was maintained within 2 mm and 2° of the stimulation target following the neuronavigation software (NBS, Nexstim Plc).

### Data pre-processing

After the data were divided into epochs and baseline-corrected, the TMS pulse artifact was removed by replacing the signal in time interval [−2, 7] ms with a cubic-spline-interpolation. Then, trials were visually inspected, and those highly contaminated by continuous muscle activity, large electrode drifts, or extensive ocular artifacts were discarded. Next, blink- and eye-movement artefacts were removed with independent component analysis (ICA). After another baseline correction and a linear detrending, the remaining scalp muscle artifact, if any, was removed with SSP–SIR from the time interval [−5, 15] ms. One more round of discarding contaminated trials by visual inspection was performed, after which the data were transformed to average reference and filtered (1–80-Hz passband and 48–52-Hz notch FIR filters).

### Data analysis

TEP data were averaged over montages of electrodes under the stimulation site. For M1, the montage contained C1, C3, CP1, CP3, FC1, and FC3. For pre-SMA, it contained F1, FC1, and FCz.

To determine a *TEP threshold*, we developed two methods based on power spectral density (PSD) analysis, named *piecewise-linear PSD fit* and *sigmoid PSD fit*, and compared their results to a fixed-amplitude approach. In the fixed-amplitude method, we calculated the TEP deflection range in the time interval [15, 50] ms post-pulse from the TEPs averaged across trials. The TEP threshold was defined as the intensity at which the deflection magnitude exceeded 6 μV (Casarotto et al., 2022; Tervo et al., 2022; Ukharova et al., 2025). The PSD-fit approaches were based on characterizing the shape of the EEG response via frequency-domain analysis and fitting sigmoid functions and piecewise linear curves into the PSD components of individual frequency bands as a function of intensity. To make the analysis generalizable between different subjects and brain regions, the methods were designed to be agnostic to specific peak timings.

PSDs of 400-ms segments of EEG data were calculated before and around the TMS pulse, yielding a frequency resolution of 2.5 Hz. The pre-pulse baseline EEG was taken from the time interval [−800, −400] ms, and the around-pulse TEP data from [−150, 250] ms. To reduce spectral leakage, the EEG data segments were windowed with a Hann window. We then baseline-corrected the PSDs by subtracting their corresponding baseline PSDs. For each stimulation condition, the PSDs were calculated from the EEG averaged across trials.

The TEP threshold in the PSD fit methods was determined by selecting the 19 PSD components from 5 to 45 Hz and determining a component threshold and a weight parameter for each. We fitted two-segment piecewise linear or sigmoid curves into the PSD component magnitudes as a function of intensity. The piecewise fit was restricted to only non-negative values and slopes to prevent unnatural results. The slope parameter of the sigmoid fit had a lower bound of 0.1 to prevent unnatural intensity ranges when fitting to near-linear datasets.

For the piecewise linear approach, the intensity at the piecewise joint was selected as the component threshold. For the sigmoid fit, in order to represent the onset of nonlinearity in the fit, we selected the intensity at the point where the sigmoid reaches 5% of its normalized maximum. The weight parameter 𝜔 was defined via the relation of the errors of the chosen fit 𝜀_alt_ and a linear fit 𝜀_lin_ such that

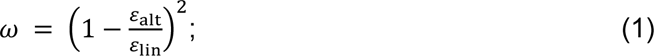

The better the nonlinear fit explains the data compared to a linear fit, the more clearly there exists a threshold where the intensity behavior changes, and thus, more weight is assigned. Consequently, the TEP threshold was defined as the weighted average of the 19 component thresholds of individual frequencies. This method is illustrated in Fig. 1.

**Fig. 1.**
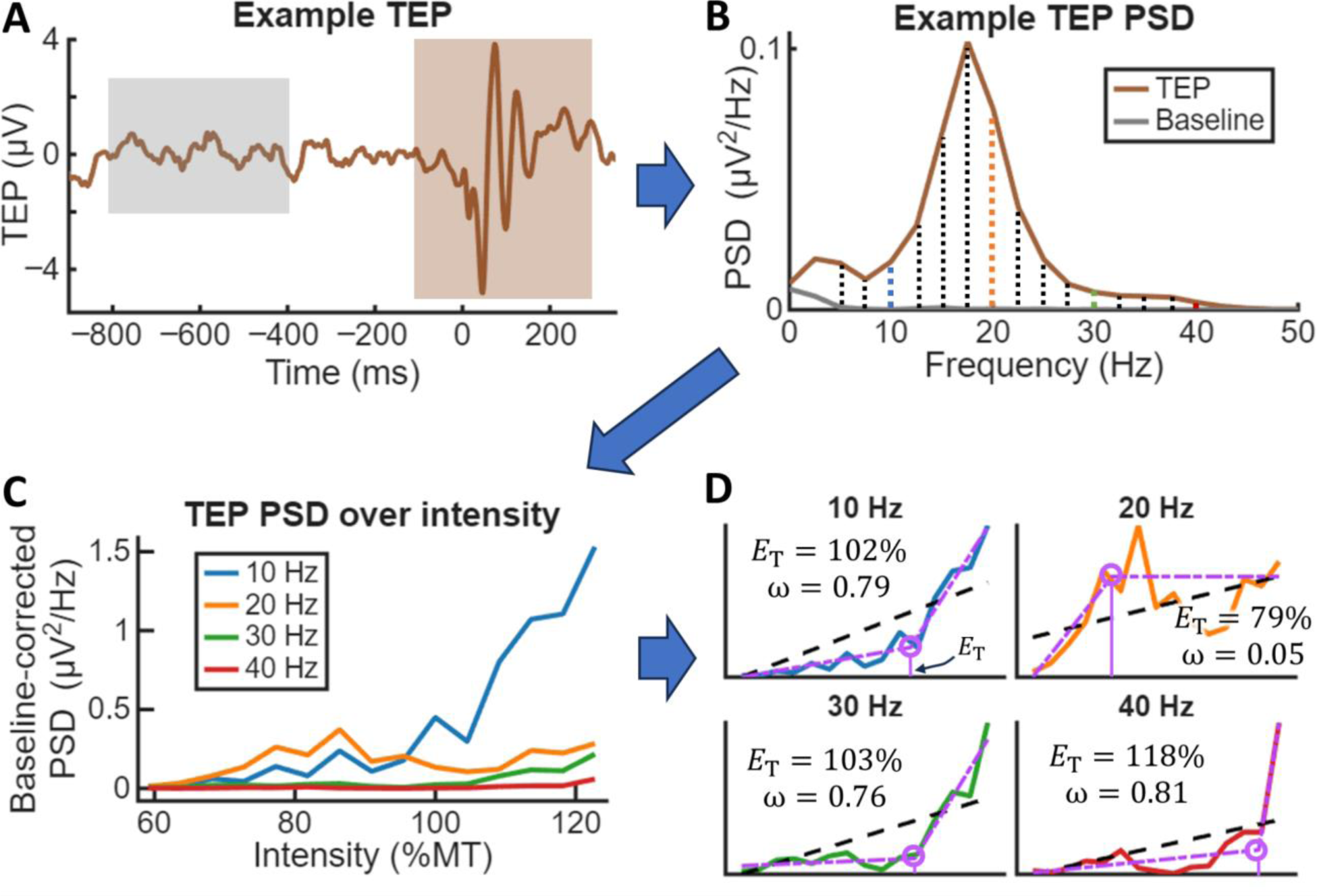
EEG-based intensity thresholding via baseline-corrected PSD. **A)** An example TEP from which we calculate PSDs in 400-ms segments both before and around the TMS pulse (*t* = 0 s). **B)** The differences between pulse- and baseline PSD in 19 frequency bands of interest. Frequencies 10, 20, 30, and 40 Hz are highlighted with different colors. **C)** Calculating the PSD at a range of intensities determines the intensity behavior of these baseline-corrected PSD components. For clarity, only the highlighted frequencies are displayed. **D)** A linear fit and either a piecewise-linear or sigmoid curve is fit to the data (piecewise fit shown). For each frequency, a component threshold 𝐸*_T_* is determined as the intensity at the joint of the nonlinear fit (circled), alongside a weight parameter 𝜔 based on the relative error between the fits (see Eq. 1). A weighted sum of these component thresholds yields the *TEP threshold*.

### Statistical analysis

We compared the results from the three TEP thresholding methods to check whether they yield similar results. In addition, we compared the results between the two targets to confirm whether the stimulation target has a statistically significant effect on the resulting TEP thresholds. To examine the effects of these factors, we applied a Scheirer–Ray–Hare test at a significance threshold of 0.05 as a non-parametric equivalent of a two-way ANOVA. If this test indicated statistically significant differences between the methods, additional Wilcoxon signed-rank tests were performed as post-hoc pairwise comparisons to explore which methods differ significantly from each other. We performed these post-hoc tests individually for both targets, resulting in six pairwise comparisons; thus, the Bonferroni-corrected significance threshold for the post-hoc tests was set at 0.0082.

## Results

A low-artifact stimulation target within M1 and pre-SMA was found for all subjects, yielding clear TEPs from 10 ms onwards. Sample TEPs from four subjects are shown in Fig. 2. The TEP-onset intensities used as references for our studied intensity ranges were found at 50–73 %MT (mean 63 %MT, SD 7.2 %MT) over M1. In pre-SMA, excluding subject 1 who had an unexpectedly high reference intensity of 98 %MT, the reference intensity was in the range of 66–86 %MT (mean 78 %MT, SD 7.6 %MT).

**Fig. 2.**
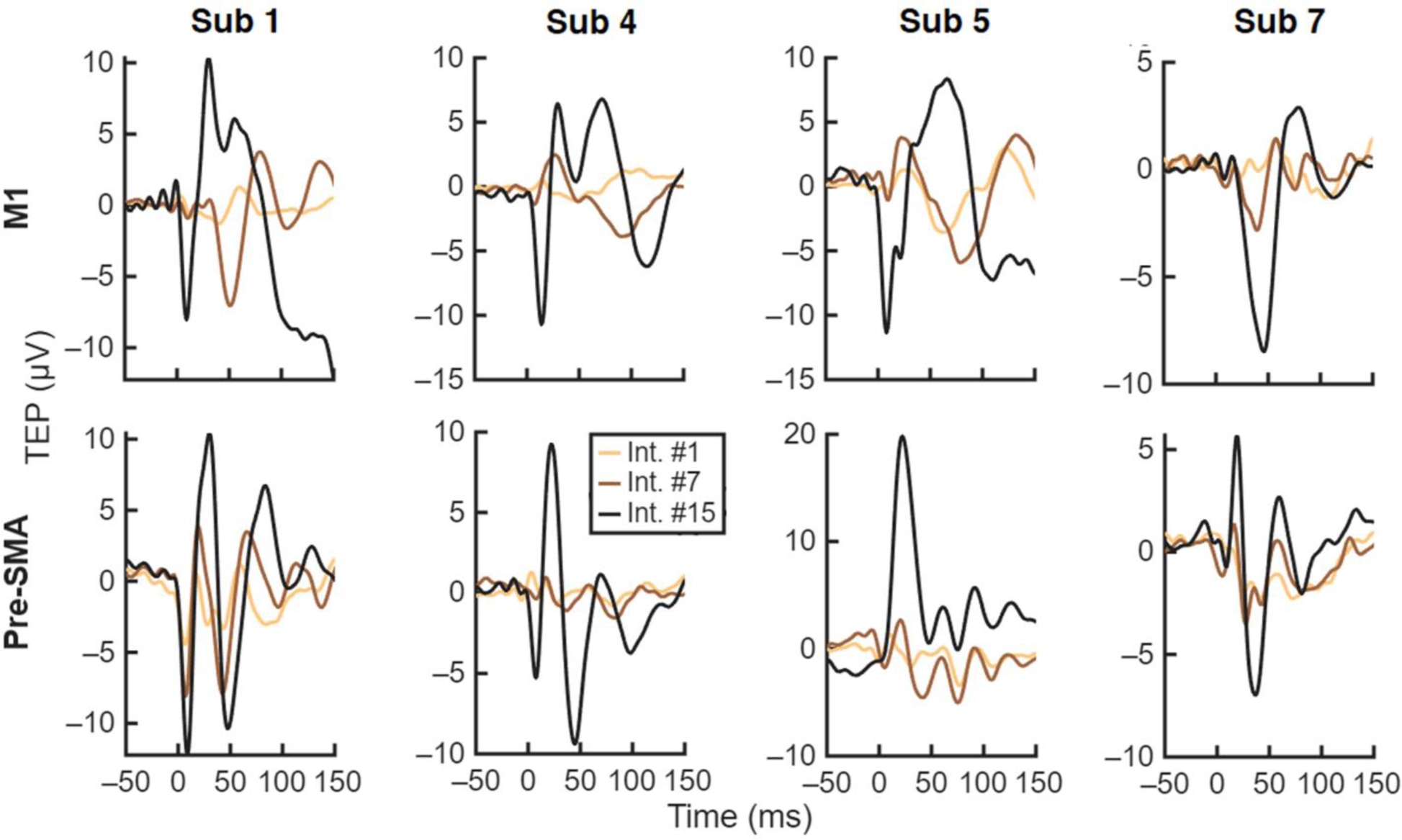
Sample TEPs averaged over roughly 60 trials from four representative subjects at both stimulation sites, at three stimulation intensities ranging from lowest to highest.

TEP thresholds were estimated for all subjects at both targets using each of the thresholding methods. The resulting threshold intensities are shown in Fig. 3. The different methods yielded significantly different TEP thresholds (*p*=0.014). The fixed-amplitude method generally yielded the lowest TEP thresholds, with the piecewise-linear method almost always resulting in higher values (M1 *p*=0.039; pre-SMA *p*=0.11), and the sigmoid method consistently yielded higher thresholds than the piecewise-linear method (M1 *p*=0.0078; pre-SMA *p*=0.0078). Consequently, the sigmoid approach always resulted in higher thresholds than the fixed-amplitude method (M1 *p*=0.0078; pre-SMA *p*=0.0078). The piecewise method yielded the most consistent results across subjects (SD 11.9 %MT at M1, 15.7 %MT at pre-SMA), followed by the fixed-amplitude threshold (SD 14.7 %MT at M1, 18.6 %MT at pre-SMA), and then the sigmoid approach (SD 17.4 %MT at M1, 22.5 %MT at pre-SMA).

**Fig. 3.**
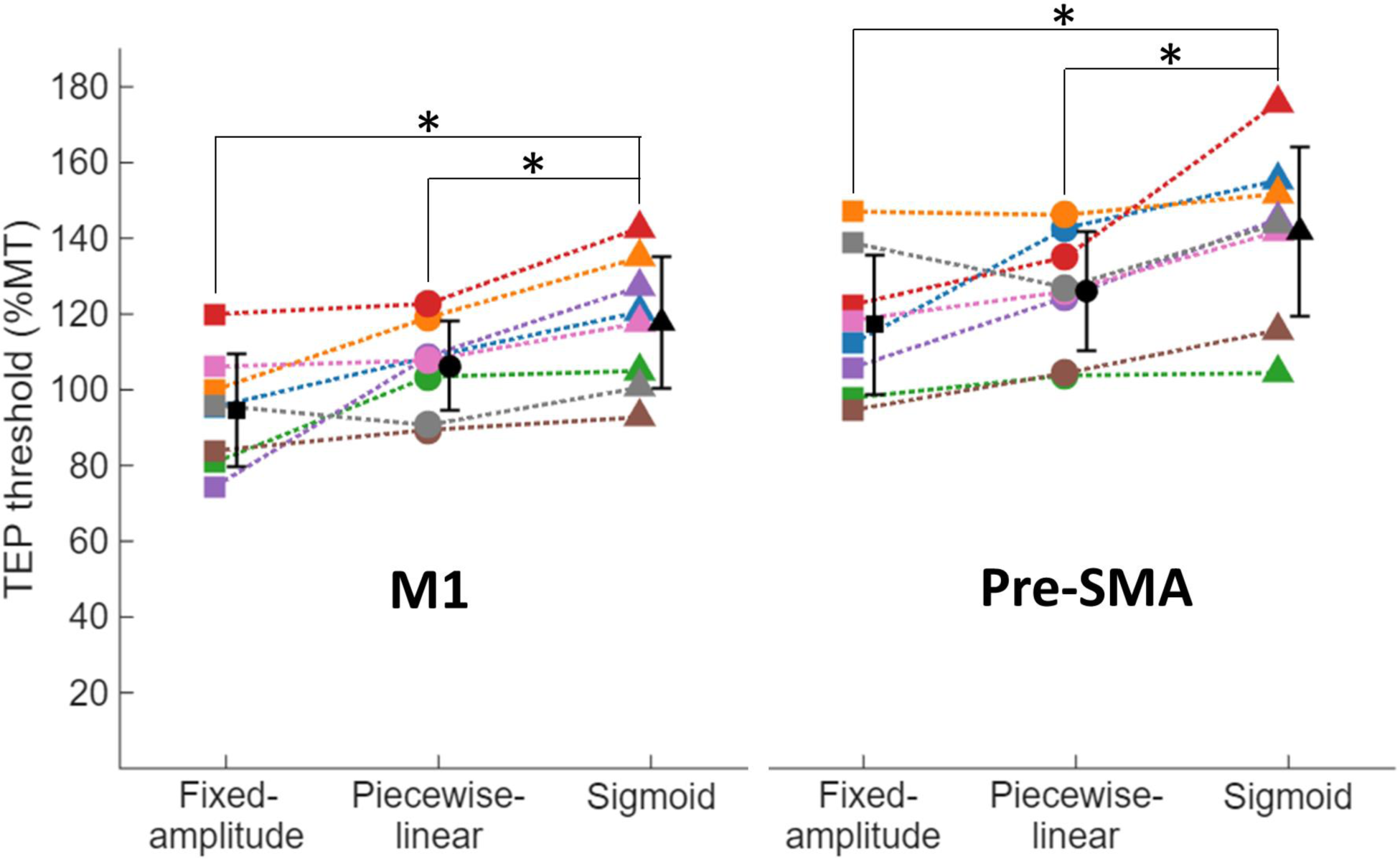
TEP threshold intensities in M1 and pre-SMA with respect to MT, as estimated by three different methods: intensity where early TEP peak-to-peak amplitude exceeds 6 µV (“fixed-amplitude”), and approaches based on fitting curves into PSD components (“piecewise-linear” and “sigmoid”). The TEP thresholds from the fixed-amplitude and piecewise-linear approaches averaged around 100 %MT in M1 and 120 %MT in pre-SMA, whereas the sigmoid method yielded higher threshold intensities. The black symbols indicate the mean and the error bars the standard deviation.

Across all methods, the threshold intensities were overall higher in pre-SMA than in M1 (*p*=0.0014). In M1, the mean threshold intensity across subjects was close to the MT at 95 and 106 %MT with the fixed-amplitude and piecewise methods, respectively. The sigmoid method resulted in a mean intensity of 118 %MT. In pre-SMA, the fixed-amplitude and piecewise methods yielded mean results close to 120 %MT (117% and 126%, respectively) and the sigmoid threshold a mean of 142 %MT.

Almost all subjects showed, at both targets, frequency ranges with low (<0.05) and high (>0.7) values of the weight parameter *ω*, indicating the presence of highly linear and highly nonlinear intensity behavior, respectively. Which frequencies displayed nonlinearity and which did not were subject-dependent. For example, in pre-SMA, the *ω* distribution of subjects 1 and 2 showed nonlinearity in only one frequency band, subjects 5 and 7 showed it in every frequency except one narrow band, subject 8 had a bifurcated distribution with frequency bands of linear or nonlinear behavior, and other subjects had broader distributions with less clear differences between frequencies. Sample distributions of *ω* from three representative subjects are shown in Fig. 4.

**Fig. 4.**
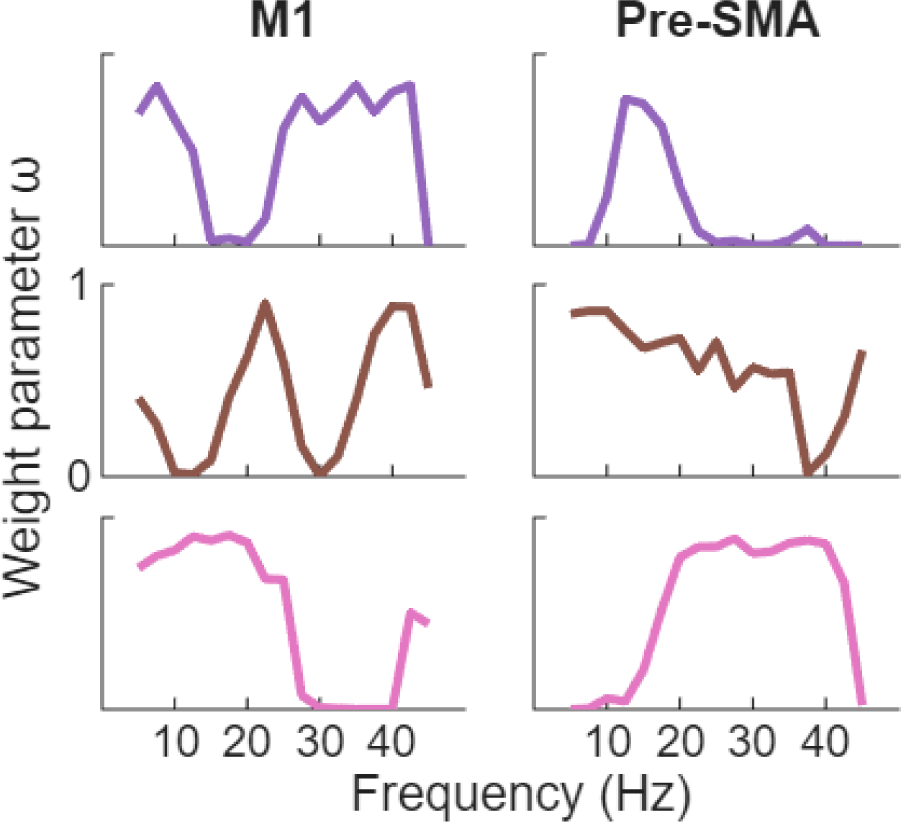
Distributions of the weight parameter *ω* at different frequencies among a sample of three subjects, determined via the piecewise method. The higher the *ω*, the more nonlinear the behavior of spectral power with respect to intensity at that frequency. The frequency bands that show nonlinear behavior are highly subject-dependent.

## Discussion

We developed and tested three methods for finding stimulation intensity thresholds for qualitative changes in EEG responses: a fixed-amplitude approach based on TEP deflection magnitude, and two PSD-fit approaches based on identifying nonlinear behavior in PSD component magnitudes of EEG data as a function of intensity. We validated these methods by identifying TEP thresholds at M1 and pre-SMA targeting sites that were artifact-free even at high intensities, using 15 closely-spaced intensities starting from the initial onset of TEPs. In M1, the fixed-amplitude and piecewise methods yielded results that, averaged across subjects, corresponded to the APB finger muscle MT, an intensity associated with functional activation. In pre-SMA, the average threshold was found at 120 %MT, showing that differences in coil–cortex distance or cytoarchitecture greatly affect the required stimulation intensities for perturbing different neuronal populations (Casarotto et al., 2010; Casula et al., 2022; Rosanova et al., 2009; Song et al., 2024).

The TEP threshold intensities varied greatly between subjects, with differences of at least 33 %MT, depending on the method and target. This highlights the importance of individualizing stimulation intensity to achieve a desired response. Intensity thresholding based on cortical responses, such as TEPs, rather than corticospinal responses, such as MEPs, can be of particular interest when attempting to elicit reliable cortical responses from cytoarchitecturally different brain areas (Ozdemir et al., 2021; Rosanova et al., 2009; Song et al., 2024). Searching for such thresholds could guide the selection of appropriate stimulation intensities for different brain circuits, and thus different treatment purposes (Chung et al., 2018; Lee et al., 2021; Lioumis et al., 2025; Tremblay et al., 2019). It can be assumed that TMS at an intensity that effectively engages the stimulation target would be more effective for treatment compared to an intensity that does not (Alhourani et al., 2015; Motzkin et al., 2023; Sack et al., 2024). The lack of such individualization may partially explain the low remission rate of some rTMS treatment interventions (Ferrarelli & Phillips, 2021; Lefaucheur et al., 2020; Lioumis et al., 2025; Parmigiani et al., 2025); a standardized intensity such as 120 %MT may not be sufficient to induce a functional brain response in all patients.

EEG-based thresholding could also be utilized to search for the minimum intensity required to engage a brain network rather than a single cortical target, as a potentially more effective way to select stimulation intensities when treating network disorders such as major depressive disorder or addiction (Lioumis et al., 2025; Wu et al., 2024). However, eliciting a TEP alone does not necessarily imply that the stimulation engages a desired brain circuit. For example, stimulating a motor hotspot in M1 can yield clear TEPs even when the stimulation does not elicit any motor responses. As stimulation intensity increases and the brain response shifts to include new functional activation, new neuronal populations and pathways are activated; thus, the shape of the EEG signal is expected to change (Casali et al., 2010; Lioumis et al., 2025; Raffin et al., 2020; Rosanova et al., 2009). Our results demonstrate this effect, as the recorded TEPs showed frequency components that increased sharply only above specific threshold intensities. These TEP thresholds could thus be more meaningful for identifying function-specific brain circuit engagement than the threshold of initial TEP onset.

Many studies have mapped various features of intensity behavior in the TMS–EEG response, from peak magnitudes of individual deflections to behavior of derived quantities such as significant current density (Casali et al., 2010; Kähkönen et al., 2005; Komssi et al., 2004, 2007; Raffin et al., 2020; Saari et al., 2018). However, these studies have generally applied only a small number of intensities across a wide range, without attempting to identify threshold intensities signifying changes in the TMS–EEG response beyond its initial onset. In contrast to our data, Kähkönen et al. (2005) found peak amplitudes of five major deflections at 60, 80, 100 and 120 %MT to exhibit linear behavior, whereas Komssi et al. (2004) reported either linear or early-sigmoid behavior in GMFA peaks at the same intensities. Raffin et al. (2020) studied TEP input–output properties at five evenly spaced intensities from 40 to 120 %MT, assessing local cortical excitability by comparing the quality of TEP linear regressions across intensities. They reported nonlinear effects in individual TEP peaks, such as P30 in M1 at 100 %MT, while other peaks remained linear, which is in line with our findings of a changing shape of the M1 TEP at around 100 %MT. In the frequency domain, Saari et al. (2018) reported linear behavior in the early components of inter-trial coherence and event-related spectral perturbation (ERSP) at the same four intensities as Komssi and Kähkönen above, in data averaged across subjects. However, similar to our data, they reported significant inter-subject differences in ERSP; averaging across subjects with nonlinear intensity behavior at different TEP thresholds may hide the nonlinear nature of the intensity dependence, explaining the difference between our results. In source space, Casali et al. (2010) looked at significant current density, phase-locking, and significant current scattering at a range of eight intensities chosen with respect to the initial onset of TEPs. While nonlinear behavior was reported in various parameters, outside Raffin et al. (2020), these works did not characterize the intensities at which such behavior occurs. Furthermore, other than Casali et al. (2010), the low intensity resolutions in the studies would make it difficult to accurately identify such thresholds.

Much like the TEP waveforms (Fig. 2), the distributions of the weight parameter *ω*, which describes nonlinearity in PSD-component intensity behavior, varied greatly between subjects (Fig. 4). This could reflect small differences in exact stimulation targets between subjects, and therefore, which brain circuits were engaged, as broader brain regions can consist of multiple smaller functional subregions (Gogulski et al., 2024; Lioumis et al., 2025). Alternatively, these differences could be explained by individual brain anatomy and structure, as TEPs are known to be subject-dependent even when observing similar functional readouts such as MEPs (ter Braack et al., 2019). These observations are relevant especially for automating TEP-based thresholding techniques.

As local cytoarchitectural differences may be reflected in the frequency ranges displaying nonlinear behavior, the distributions of *ω* could be exploited not only to determine the appropriate stimulation intensity but also to identify the target. In the future, a real-time readout of PSD may be utilized as a marker of perturbation of different neuronal populations, supplementing demanding neuroimaging-guided TMS–EEG mapping approaches (Ukharova et al., 2025) for selecting an appropriate stimulation location. However, further research is required to explore the relationship between inter-subject differences in *ω* distributions and the stimulated circuits.

Our approach considered only the electrodes near the stimulation site. A more sophisticated EEG-based identification of intensity thresholds could involve studying the TMS-induced signal propagation across the cortex. Previous research has suggested that low-intensity TMS results in a mostly local perturbation, whereas stronger stimulation may engage brain networks that carry the signal to other brain regions (Casali et al., 2010; Garcia et al., 2011). As many brain disorders are understood to be network disorders, identifying the individualized intensity required to induce particular network activation could be of significant clinical benefit. However, designing metrics relying on connectivity estimates may require more prior knowledge to be effective, such as subject- and stimulation target-dependent information on where the signal is expected to propagate, rendering such methods more complicated than the local excitability estimates introduced here.

The measurements were performed with each condition in a separate block due to software limitations. As the state of the subject and the position of the coil may differ slightly between blocks, extra variability may have been introduced to the data. This noise could have been eliminated by randomizing measurement conditions within each measurement block. We expect such a measurement would yield clearer intensity behaviour in PSD components, and thus more reliable TEP thresholds. Additionally, while our intensity mapping aimed to find the highest intensity without a TEP, our starting intensities generally yielded clearly observable TEPs when averaged over the full 60 trials, suggesting that 20 trials is not enough to distinguish early, low-amplitude TEPs from baseline. While the true earliest peaks likely occurred at slightly lower intensities, this does not compromise our results, as the relevant EEG behavior was still captured within our intensity range.

## Conclusion

We demonstrated that the waveform of the TMS-evoked EEG response, characterized by its PSD, changes at particular intensity thresholds when stimulating M1 and pre-SMA. These TEP thresholds vary between subjects and cortical areas, and they may signify a qualitative shift in brain response, such as a new brain network becoming engaged. Thus, selecting the stimulation intensity based on TEP thresholds could help ensure that the stimulation strength is sufficient to induce a desired brain activation. Individualizing the stimulation intensity may significantly improve the efficacy of TMS as a treatment compared to existing protocols that use predetermined percentages of the motor threshold for all subjects.

