## Supplementary material for "TMS–EEG Indices to Define Local Cortical Excitability Thresholds"

**Individual TEPs**

60 TMS–EEG trials were recorded from eight subjects at 15 closely spaced intensities at both M1 and pre-SMA, starting from the estimated intensity for the initial onset of a visible TEP. Traces of these measurements are shown in Fig. 1 (M1) and Fig. 2 (pre-SMA).

While many of the subjects showed similar overall TEP features when stimulated at the same brain region, high inter-individual variability was also observable. For example, the responses to M1 stimulation in subjects 3 and 7 showed fewer peaks than in other subjects. As for intensity-induced changes in TEP shape, in M1, a sudden shift in TEP behavior from 50 ms onwards occurred at high intensities in most subjects. In pre-SMA, the changes were less pronounced, but some subjects displayed TEP behavior that only occurred at high intensities, such as a peak emerging at roughly 30 ms in subject 1 only with sufficiently high intensities.

TEPs are often characterized by the amplitudes and latencies of individual peaks. However, we found that the presence and/or latencies of the peaks varied between intensities: some peaks stayed constant in time while others consistently increased in latency, and some peaks were subsumed into neighboring peaks as intensity increased.

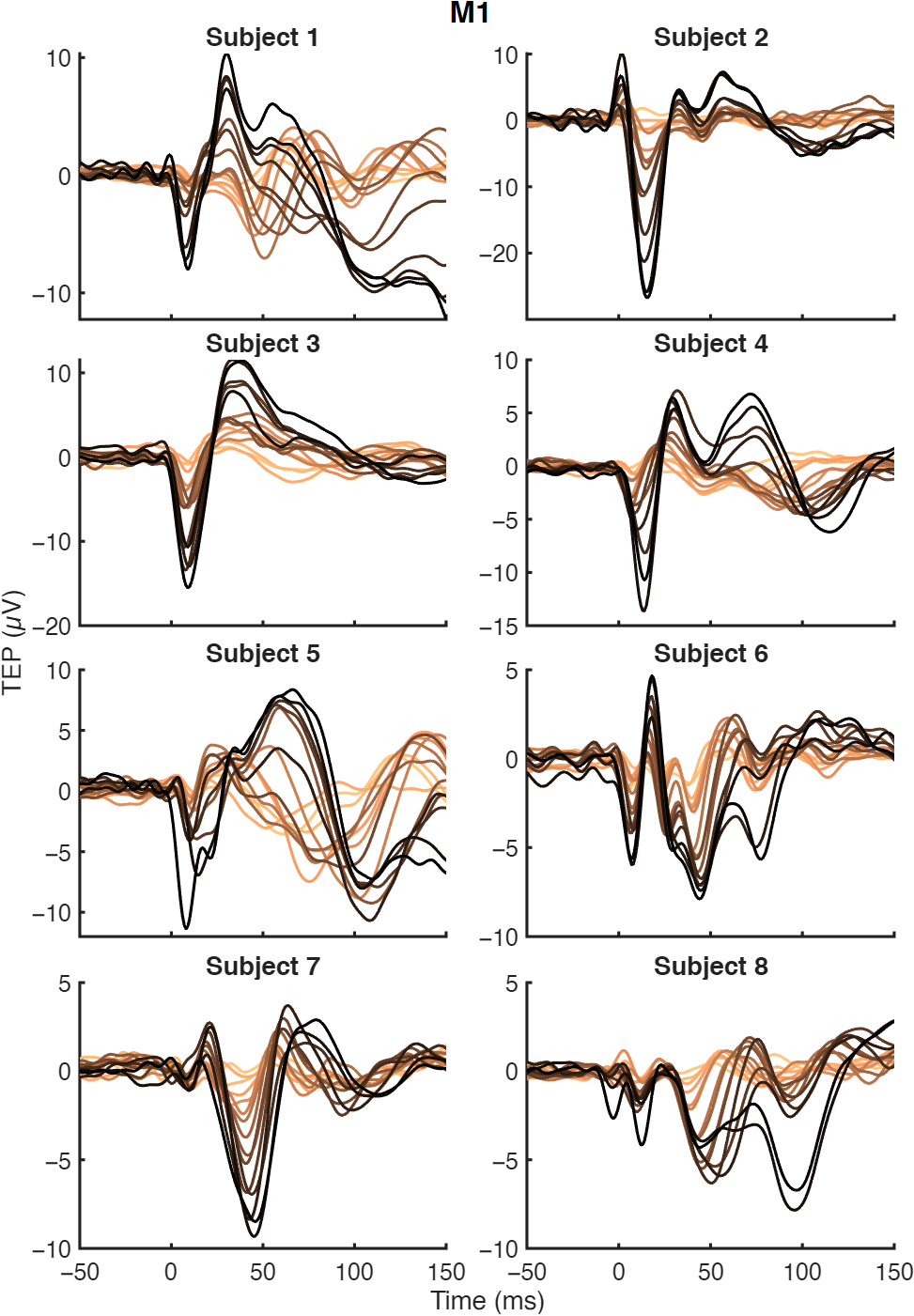
Fig. 1. The full range of TEP traces elicited from M1 by the 15 stimulation intensities, darker colors corresponding to stronger stimulation.

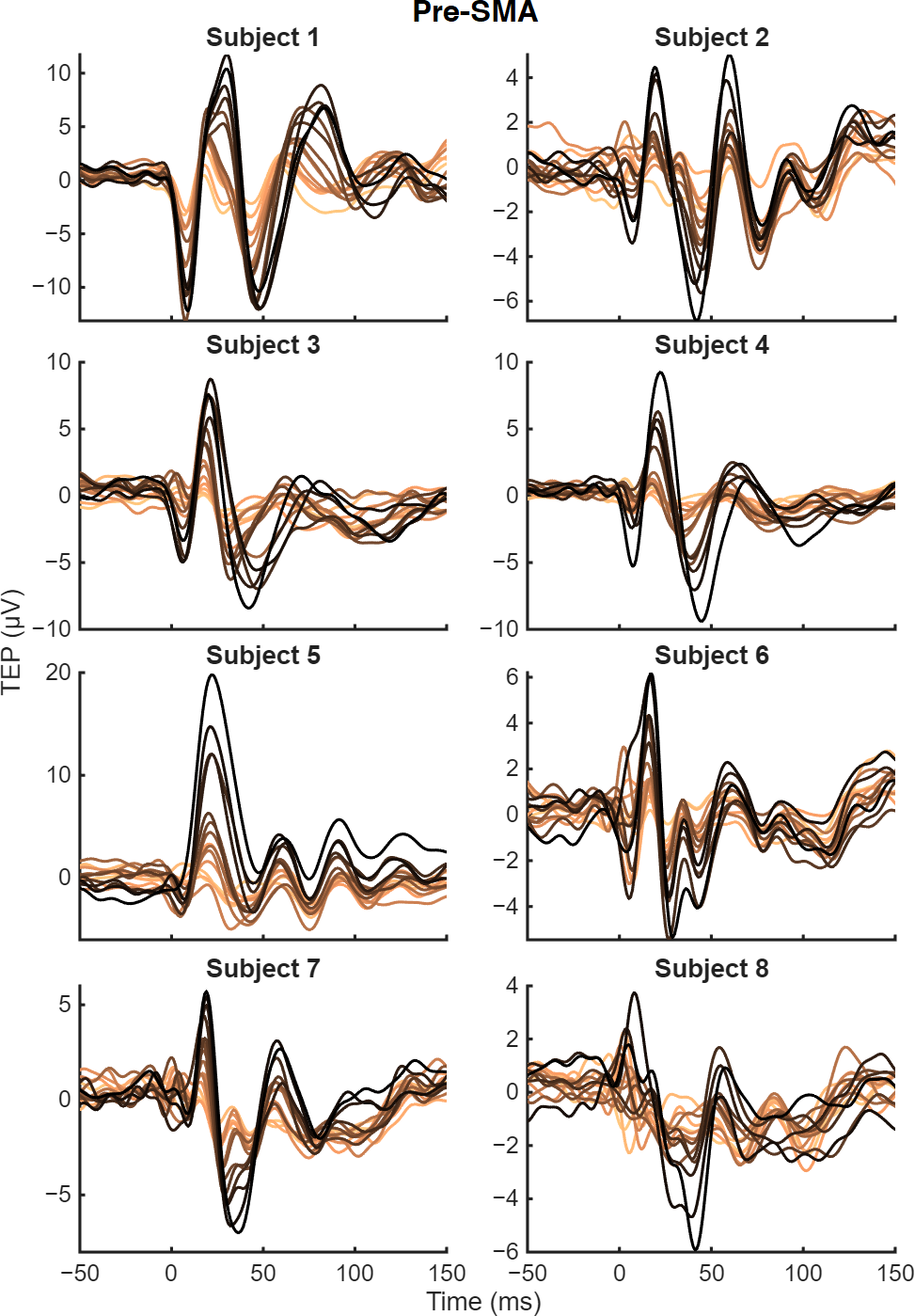
Fig. 2. The full range of TEP traces elicited from pre-SMA by the 15 stimulation intensities, darker colors corresponding to stronger stimulation.

**Intensity thresholds and weight parameter** $\boldsymbol{\omega}$

For each subject and target, the intensity behavior of individual frequency components was studied by calculating the power spectral density (PSD) of a 400-ms-segment of the EEG around the TEP. We searched for nonlinearity in the PSD intensity behavior of the frequency components from 5 to 45 Hz, assigning each a threshold intensity indicating at what intensity the behavior changed, and a weight parameter $\omega$ that quantified the nonlinearity of the data. The TEP threshold was then defined as the $\omega$-weighted mean of the individual intensity thresholds; frequency bands exhibiting clearer nonlinearity were considered more relevant. The distributions of the individual intensity thresholds and the weight parameters $\omega$ for each subject at both targets are shown in Fig. 3. The data in this figure were obtained with the piecewise-linear PSD fit method for threshold identification.

In almost all cases, $\omega$ strongly depended on the frequency, with these distributions being highly variable between subjects and targets. The threshold intensity, however, was often nearly constant across frequencies, with variation usually corresponding to large shifts in $\omega$, such as with subject 1 at pre-SMA. This implies that when clear nonlinearity is present in the intensity behavior of the PSD component, the thresholds are largely invariant with respect to frequency, but when the nonlinearity is ambiguous, estimates for where the shift in behavior occurs become unreliable. However, in some subjects, different frequency bands with high $\omega$ had slightly different thresholds. For example, subject 1 in M1 showed thresholds at around 100 %MT in the low-frequency components, whereas high-frequency components had thresholds at around 120 %MT. This could suggest that different cell types, neuronal populations or circuits start contributing to the recorded signals at different intensities, manifesting as distinct EEG response characteristics.

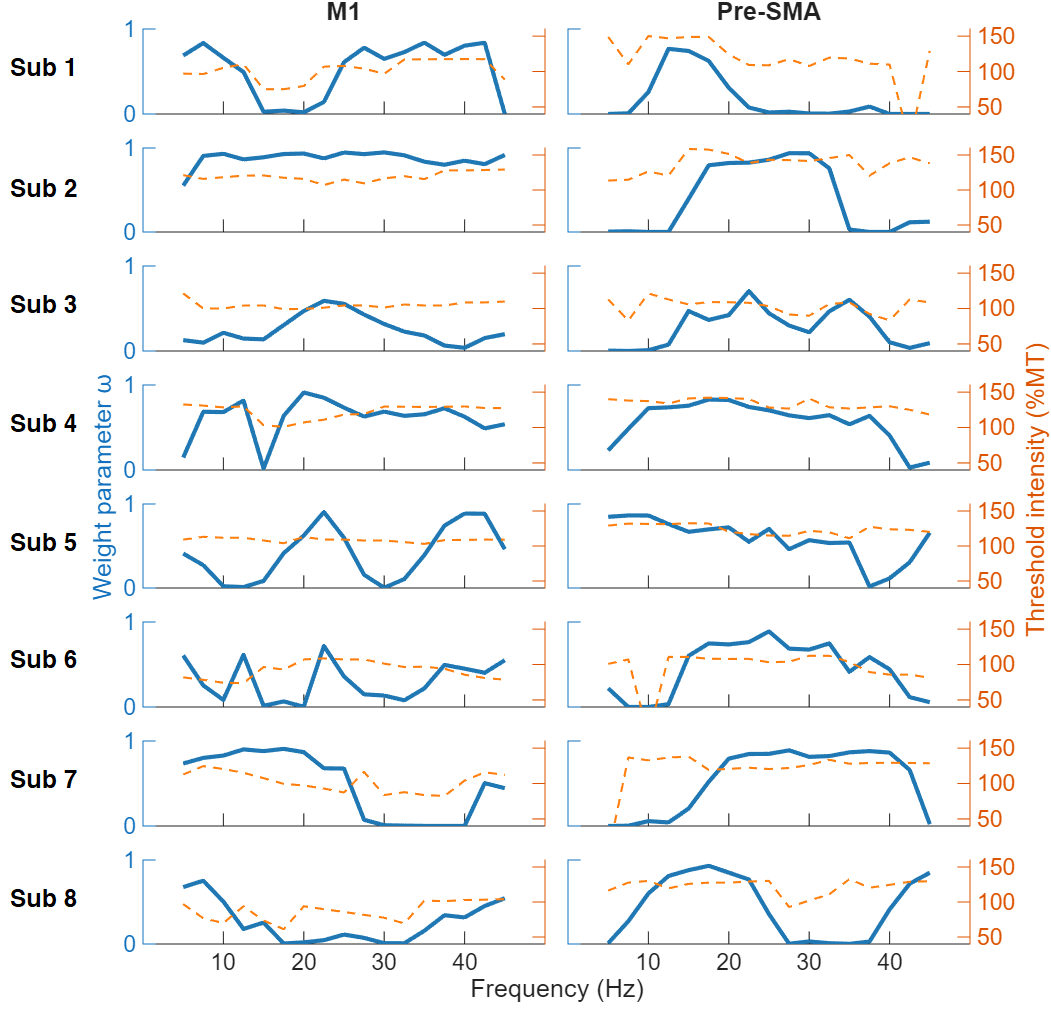

Fig. 3. The frequency distributions of weight parameter $\omega$, as well as the frequency-component-specific intensity thresholds for all eight subjects. The intensity thresholds were defined as the points of nonlinearity as determined by the piecewise-linear PSD fit method.
